# LRRK2 kinase activity restricts NRF2-dependent mitochondrial protection in microglia

**DOI:** 10.1101/2024.07.09.602769

**Authors:** Chi G. Weindel, Lily M. Ellzey, Aja K. Coleman, Kristin L. Patrick, Robert O. Watson

**Author notes:** Correspondence Twitter: @The_PW_Lab.

## Abstract

Mounting evidence supports a critical role for central nervous system (CNS) glial cells in neuroinflammation and neurodegenerative diseases, including Alzheimer’s disease (AD), Parkinson’s Disease (PD), Multiple Sclerosis (MS), as well as neurovascular ischemic stroke. Previously, we found that loss of the PD-associated gene leucine-rich repeat kinase 2 (*Lrrk2*) in macrophages, peripheral innate immune cells, induced mitochondrial stress and elevated basal expression of type I interferon (IFN) stimulated genes (ISGs) due to chronic mitochondrial DNA engagement with the cGAS/STING DNA sensing pathway. Here, we report that loss of LRRK2 results in a paradoxical response in microglial cells, a CNS-specific macrophage population. In primary murine microglia and microglial cell lines, loss of *Lrrk2* reduces tonic IFN signaling leading to a reduction in ISG expression. Consistent with reduced type I IFN, mitochondria from *Lrrk2* KO microglia are protected from stress and have elevated metabolism. These protective phenotypes involve upregulation of NRF2, an important transcription factor in the response to oxidative stress and are restricted by LRRK2 kinase activity. Collectively, these findings illustrate a dichotomous role for LRRK2 within different immune cell populations and give insight into the fundamental differences between immune regulation in the CNS and the periphery.

## INTRODUCTION

Microglia are slow dividing long-lived resident macrophage populations of the CNS (1, 2), that develop from yolk sac myeloid hematopoietic precursors and travel to the brain prior to closure of the blood brain barrier (3–6). As an innate immune population, microglia act as the first line of defense against invading pathogens and cellular damage through multiple processes including phagocytosis, neuron pruning, cytokine release, and direct cell-cell communication (7–12). The tight regulation of these processes is necessary to prevent neuroinflammation, a hallmark of neurodegenerative diseases of the CNS (13–16).

The type I IFN response has gained recent interest for its role in both maintaining neuron health as well as promoting inflammation. Signaling through the IFNα/β receptor (IFNAR) has been shown to be critical for the development of healthy neurons, as cell-type specific loss of IFNAR signaling results in Lewy body accumulation, α-synuclein aggregation, and a PD-like phenotype due to a blockade of neuronal autophagy (17). Microglial type I IFN responses also play crucial roles in neurodevelopment, where IFNAR signaling facilitates phagocytosis and clearance of damaged neurons (18). Although type I IFN is critical for healthy development of the CNS, IFNAR signaling can act as a double-edged sword to promote neurodegeneration. For example, in the 5xFAD model of amyloid β-induced Alzheimer’s, blocking IFNAR in CNS cell populations was overall protective against memory loss and synaptic damage (19, 20). In naturally aging brains, type I IFN signatures in microglial cells are enhanced (21) and are associated with low level inflammation, bystander cell activation, and microgliosis (22). Given what we know, increased IFN signatures could be a product of elevated microglial phagocytic activity and protection, or aberrant inflammation, bystander cell activation, and CNS damage. Thus, there is a critical need to better understand the regulatory nodes that govern protective vs. pathogenic type I IFN responses in the brain, and how this regulation drives the maintenance of healthy glia and neurons while restricting neuroinflammation.

In the peripheral immune system, the tempering of type I IFN responses is a critical means to restrict inflammation and prevent interferonopathies and autoimmunity. One well-known restrictive pathway is the NRF2-mediated redox response. Classically, NRF2 is a transcription factor that upregulates antioxidant proteins to protect against oxidative stress. NRF2 has also shown to regulate immune signaling; it is a negative regulator of type IFN during viral infection (23, 24) and can restrict IFNβ activation and inflammation following LPS stimulation or sepsis (25–27). NRF2 restricts the type I IFN response at several nodes, including inhibiting dimerization of the transcription factor IRF3 (28–30), and reducing STING expression and mRNA stability in human cells (31). Proteins upregulated by NRF2 during oxidative stress such as HMOX1 also have regulatory effects on the type I IFN response through degradation of transcription factors IRF3/IRF7 by autophagy (32), suggesting a complex interplay between the two pathways. Less is known about the connection between NRF2 and type I IFN responses in the brain. It has been shown that mice lacking NRF2 have neuroinflammation with astrogliosis (33), and NRF2 modulation impacts neuroinflammation in several PD models (34, 35). Despite the intriguing connections between NRF2 and brain health, the role of NRF2 in glial cell inflammation remains under studied.

LRRK2 is a multifunctional PD-associated kinase expressed in neurons and immune cell populations, including monocytes and macrophages (36). LRRK2 has been implicated in both genetic and sporadic forms of PD, making *Lrrk2* KO and mutation studies excellent models to investigate overall mechanisms of PD (37–39). While LRRK2 has been well studied in the context of neurons, less is known about the function of LRRK2 in immune cell populations. Previously, we found that loss of LRRK2 in peripheral macrophages including BMDMs, peritoneal macrophages, macrophage cell lines, results in significantly elevated basal levels of type I IFN/ISGs and an inability to induce interferon responses after infection (40). We linked these elevated basal I IFN responses to mitochondrial stress, namely mitochondrial fragmentation and oxidative stress, that leads to mitochondrial DNA leakage into the cytosol and chronic engagement of the cGAS/STING signaling pathway (40).

Here, motivated by our *Lrrk2* KO macrophage findings, we investigated how loss of LRRK2 impacts CNS resident microglia cells. Surprisingly, we found that microglial cells lacking LRRK2 had a reduction in type I IFN transcripts compared to controls. Differential gene expression analysis uncovered that the NRF2 redox pathway was upregulated in *Lrrk2* KO microglia. Consistent with enhanced protection, *Lrrk2* KO microglial cells were better at maintaining a healthy mitochondrial membrane potential and a higher capacity for OXPHOS, which was dependent on NRF2. Finally, inhibition of LRRK2 kinase activity was sufficient to reduce ISGs and upregulate NRF2, indicating that LRRK2 kinase function plays an active role in regulating the type I IFN response and NRF2 in microglial cells. This work gives insight into the functional requirements of different macrophage populations and how anti- and pro-inflammatory processes are regulated in various tissues.

## RESULTS

### The type I IFN signature is reduced in microglial cells upon loss of LRRK2

To understand how loss of LRRK2 impacts microglial cells, we first developed a process to generate and isolate pure populations of non-activated primary microglial cells from the cerebral cortices of P1.0-P1.5 neonates using sex and age matched littermate controls (*Lrrk2* KO vs. *Lrrk2* HET). Key to this approach is the ability to separate microglial cells from astrocytes, another abundant glial cell population in the brain. We achieved this by dissecting out cerebral cortices followed by a high trypsin digestion 2.5% to disrupt cells of the meninges followed by 6 days of culture for astrocytes, or 10 days of culture for microglial cells in complete media containing MCSF. Microglial cells were gently washed off the astrocyte layer with PBS/EDTA. We then verified microglial vs. astrocyte cell populations measuring GFAP mRNA expression (an abundant astrocyte transcript) and IBA1 protein (a key surface marker of microglia) (**Fig 1A**.). Astrocyte purity was also assessed by measuring GFAP+ cells (>90%) by flow cytometry (**Fig S1A**). Microglia, defined as CD45+ CD11b+, were measured to be >85% pure by flow cytometry (**Fig S1B**). To identify the major LRRK2-dependent differences in transcript abundance, we performed bulk RNA-SEQ analysis from RNA collected from resting microglia and generated sequencing libraries. Using the Rosalind RNA-seq platform, we compared transcripts between HET and *Lrrk2* KO microglia and identified 96 significantly differentially expressed genes (Adj. p-value < 0.05), with 77 downregulated and 19 upregulated genes (**Fig 1B, Fig S1B, Table S1**). Notably, many of the most significantly down regulated genes, aside from *Lrrk2*, were ISGs including *Rsad2*, *Ifit2*, *Mx1*, *Ifit3*, *Cxcl10*, *Cmpk2*, among others (**Fig 1B, 1D purple**). Consistent with the type I IFN pathway being impacted, pathway analysis through NCATS BioPlanet identified the most significant pathway impacted as IFN α/β signaling (**Fig 1C**). Significantly upregulated genes were fewer, these genes (*Id1*, *Id3*, *Dmpk*, *Cd34*, *Serpine2 and Dhfr*) have been associated with neurogenesis and microglial cell division (43–47) (**Fig 1E**). qRT-PCR confirmed downregulation of multiple ISG transcripts including but not limited to *Rsad2*, *Ifit1*, *Gbp2*, *Isg15*, and *Irf7* (**Fig 1F**). Importantly, downregulation of ISGs was also observed at the protein level (**Fig 1G, Fig S1D**). By generating lentiviral shRNA knockdowns (KDs) of *Lrrk2* alongside a scramble (SCR) control, we observed the same reduced type IFN signature in the spontaneously immortalized microglial cell line SIMA9 (**Fig 1H**), suggesting a common role for LRRK2 in promoting ISG expression in microglial cells.

**Figure 1:**
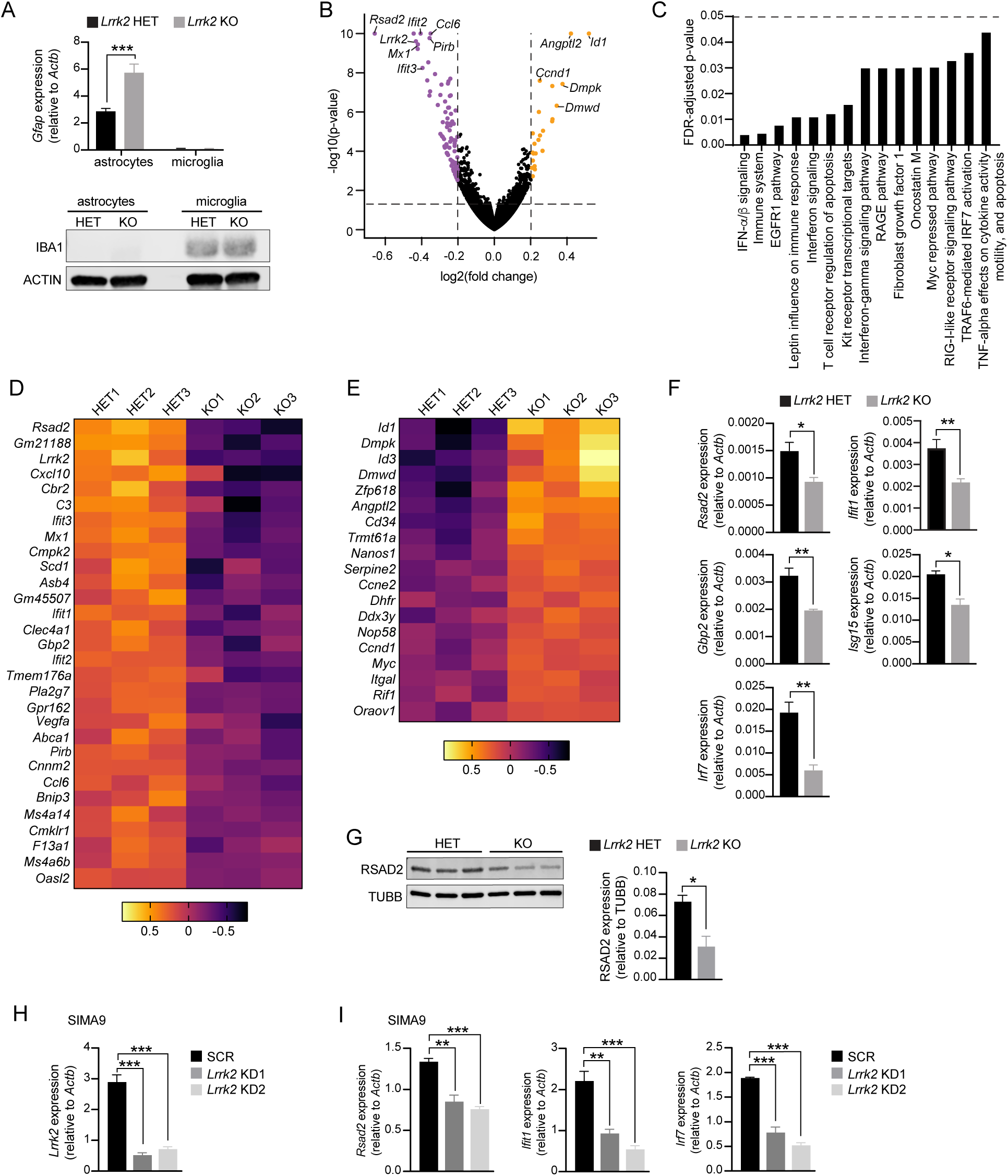
Loss of LRRK2 reduces tonic IFN signaling in microglial cells. (**A.**) Transcript levels of *Gfap* in *Lrrk2* KO and *Lrrk2* HET (control) astrocytes and microglia measured by qRT-PCR (upper graph). Protein levels of IBA1 relative to ACTIN in *Lrrk2* KO and HET astrocytes and microglia measured by western blot (lower graph). (**B.**) Volcano plot of genes differentially expressed between *Lrrk2* KO and HET microglia (left, purple) down in KO, (right, orange) up in KO. (**C.**) Ingenuity pathway analysis of major transcriptional pathways differentially expressed between *Lrrk2* KO and HET microglia (**D.**) Heatmap of significant genes downregulated in *Lrrk2* KO microglia compared to HET controls. (**E.**) Heatmap of significant genes upregulated in *Lrrk2* KO microglia compared to HET controls. (**F.**) Transcript levels of ISGs *Rsad2*, *Ifit1*, *Gbp2*, *Isg15*, and *Irf7* in *Lrrk2* KO and HET microglia measured by qRT-PCR. (**G.**) Protein levels of RSAD2, compared to TUBB in *Lrrk2* KO and HET microglia measured by western blot; n=3. Quantification on right. (**H.**) Transcript levels *Lrrk2* in *Lrrk2* KD and SCR SIMA9 microglia measured by qRT-PCR. (**I.**) The same as in (H), but ISGs *Rsad2*, *Ifit1*, and *Irf7*. Two-tailed Student’s t-test was used to determine statistical significance. *p<0.05, **p<0.01, ***p<0.005.

To better understand potential drivers of this phenotype, we looked to known regulators of the type I IFN response in macrophages (48, 49). We saw no major differences in activation (I-Ab) or costimulatory marker (CD86) expression, indicating that loss of LRRK2 did not impact major states of cellular activation (**Fig S1E**). Likewise, we saw no difference in the expression of negative regulators of type I IFN gene expression like *Sosc1*, *Smad2/3*, and *Pias4* (**Fig S1F**) (50). We also confirmed that genes associated with M1-like vs M2-like macrophage states were not differentially regulated in *Lrrk2* KO microglia, because the type I IFN response has been linked to M1 polarization through *Irf7* (51). Normal M1- and M2-like markers indicate a loss of LRRK2 and downregulation of ISGs does not play a role in polarity skewing of microglia (**Fig S1G**) Taken together, these data argue that LRRK2 is required to maintain tonic levels of ISGs in microglial cells through a previously undescribed mechanism.

### Loss of LRRK2 differentially impacts microglia and peripheral macrophages

Given the surprising phenotype of *Lrrk2* KO microglia, we decided to compare the transcriptional profile differences of *Lrrk2* KO microglia to *Lrrk2* KO peripheral macrophages. We observed an overlap of 26 genes differentially expressed genes (DEGs) in both *Lrrk2* KO microglia and macrophages, with 67 and 352 DEGs distinct for microglia and macrophages respectively (**Fig 2A, Table S1, S2**). Within the group of 26 shared DEGs, many genes were oppositely impacted by loss of LRRK2 in macrophages and microglial cells, with 19 genes increased in macrophages but reduced in microglia (**Fig 2B**). Half of these “conflicting” DEGs, including *Mx1*, *Ifit1-3*, *Oasl2*, and *Gbp2* etc., are associated with the type I IFN response and Ingenuity Pathway Analysis identified “IFNα/β Signaling” as the most significantly impacted pathway shared between *Lrrk2* KO macrophages and microglia (**Fig 2D**).

**Figure 2:**
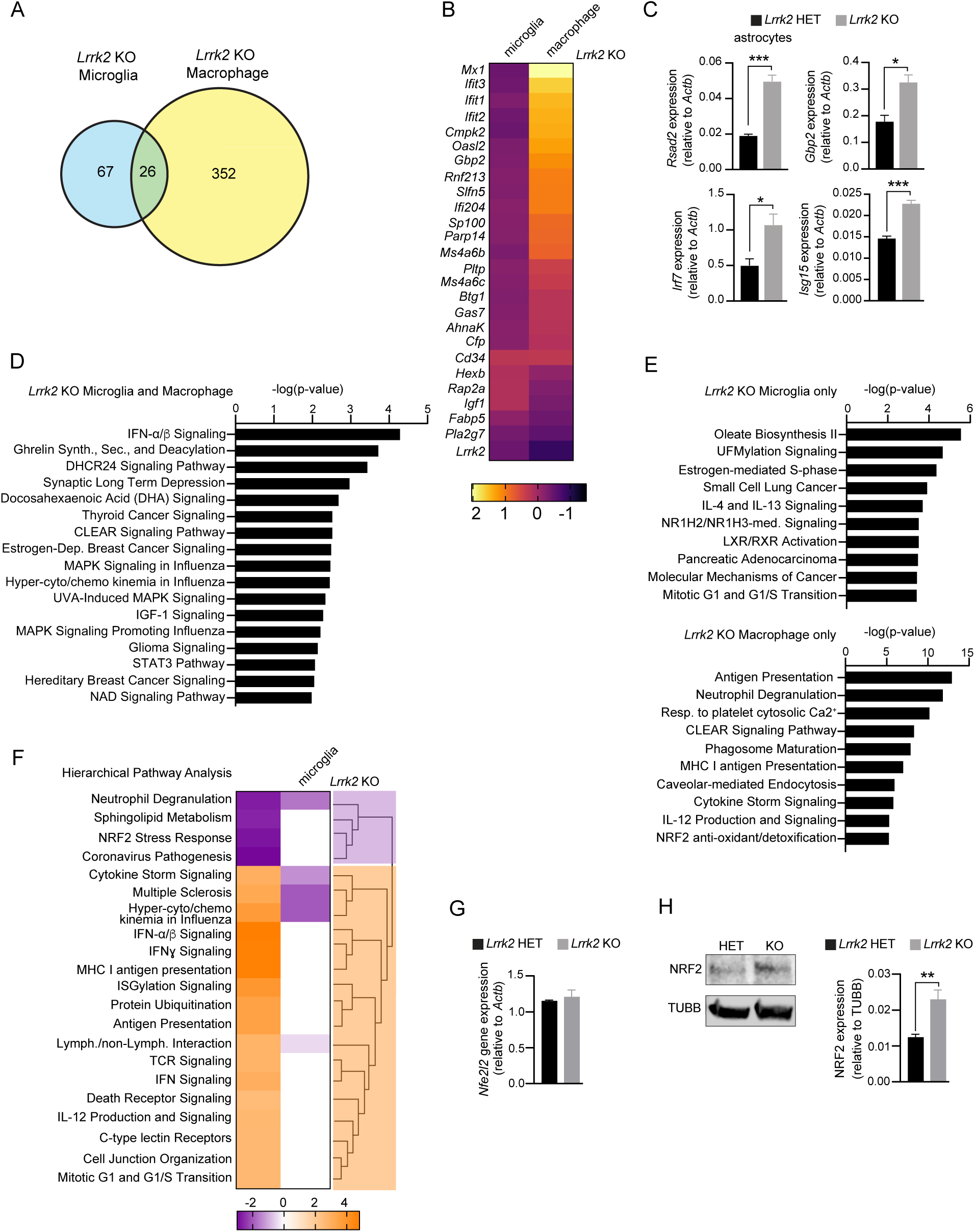
The type I IFN response and NRF2 pathways are opposing in microglial cells and macrophages. (**A.**) Venn diagram depicting genes differentially expressed in *Lrrk2* KO microglial cells (left), both *Lrrk2* KO microglia and macrophages (center) or only *Lrrk2* KO macrophages (right). (**B.**) Heatmap of genes differentially expressed in *Lrrk2* KO microglia and macrophages. (**C.**) Transcript levels of ISGs *Rsad2*, *Gbp2*, *Irf7*, *Isg15*, in *Lrrk2* KO and HET astrocytes measured by qRT-PCR. (**D.**) IPA pathway plot of shared pathways impacted by a loss of LRRK2 in microglia and macrophages. (**E.**) Pathway analysis of genes upregulated in *Lrrk2* KO microglia only (upper) and Lrrk2 KO macrophages only (lower). (**F.**) Hierarchal clustered heatmap depicting z-scores and pathway differences between microglial cells and macrophages. (**G.**) Transcript levels of *Nrf2* in *Lrrk2* KO and HET microglia measured by qRT-PCR (**H.**) The same as in (G) but NRF2 proteins levels relative to TUBB. Two-tailed Student’s t-test was used to determine statistical significance. *p<0.05, **p<0.01, ***p<0.005.

We initially hypothesized that the type I IFN phenotype in *Lrrk2* KO microglia could have something to do with the CNS residency of these cells. To test if other CNS glial cells also downregulated type I IFN in the absence of LRRK2, we performed qRT-PCR analysis on *Lrrk2* KO and HET astrocytes, purified as described in (**Fig 1A, S1A**). Unexpectedly, we found that ISGs were upregulated in *Lrrk2* KO astrocytes compared to HET controls (**Fig 2C**). We also noted that transcripts associated with astrocyte maturation and activation, e.g. *Gfap*, *S100b*, *Icam1*, and *Ccl5,* were also elevated in *Lrrk2* KO astrocytes (52) (**Fig S2**). These data suggest that *Lrrk2* KO astrocytes display a more macrophage-like type I IFN phenotype and that the downregulation of ISGs in *Lrrk2* KO microglia is not shared by other glial populations.

To better understand why loss of LRRK2 differentially impacts ISG expression in astrocytes/macrophages vs. microglia, and perhaps identify the driver of the phenotype, we performed pathway analysis of non-type I IFN genes in *Lrrk2* KO microglia and macrophages. We found that non-ISG DEGs in *Lrrk2* KO microglial cells were enriched in pathways related to cell cycle, metabolism, and cancer cells (**Fig 2E** upper graph), whereas non-ISG DEGs in *Lrrk2* KO macrophages were enriched in pathways related to in immune mediated processes, including antigen presentation, neutrophil degranulation, cytokine signaling, and NRF2-mediated antioxidant protection (**Fig 2E** lower graph). To begin to understand what differentially regulated pathways might be contributing to the inverse phenotype of *Lrrk2* KO macrophages vs. microglia, we performed hierarchical clustering of the most significant differentially regulated pathways. (**Fig 2F**). Two major clusters were noted. One cluster contained upregulated inflammatory signaling pathways (**Fig 2F**, orange box); the other contained several pathways downregulated only in *Lrrk2* KO macrophages, including sphingolipid metabolism, NRF2 stress response, and pathogenesis of coronavirus (**Fig 2F**, purple box). We chose to focus on NRF2 due to its previous association with downregulating type I IFN responses (23, 25, 32).

To establish that NRF2 expression could be impacted by loss of LRRK2, we performed qRT-PCR (**Fig 2G**), and western blot analysis (**Fig 2H**) of NRF2 in *Lrrk2* KO and HET microglia. We found that NRF2 expression was increased at the protein but not mRNA levels indicating NRF2 redox sensing is upregulated in the absence of LRRK2. Taken together, these data suggest that compared to peripheral macrophages, microglial cells employ additional regulatory nodes to restrict type I IFN responses that rely on the redox regulator NRF2.

### *Lrrk2* KO microglia are protected from stressors and have enhanced mitochondrial metabolism during activation

Because we previously found that the increase in LRRK2-dependent basal type I IFN in macrophages was linked to mitochondrial dysfunction, including reduced mitochondrial membrane potential and decreased OXPHOS (40), we hypothesized that *Lrrk2* KO microglial mitochondria would have the inverse phenotype. To test this, we performed mitochondrial membrane potential assays using JC-1 and TMRE on *Lrrk2* KO and HET microglia. JC-1 is a carbocyanine dye that accumulates in mitochondria with robust membrane potential to form red fluorescent aggregates. Upon loss of mitochondrial membrane potential JC-1 diffuses to the cytosol as a monomer where it emits a green fluorescence, providing a facile tool to determine mitochondrial health. In line with our hypothesis, loss of LRRK2 resulted in increased mitochondrial membrane potential (more red aggregates) in resting microglia and microglia exposed to rotenone/ATP (**Fig 3A**). We further confirmed this by using the mitochondrial membrane potential dye TMRE in resting *Lrrk2* KO and HET microglia (**Fig S3A, Fig 3B**), and microglia treated with the uncoupling agent FCCP (**Fig 3B**). Consistent with an inverse correlation between type I IFN upregulation and mitochondrial health, we saw the opposite response in astrocytes (**Fig S3C**). Protection of mitochondrial membrane potential was also observed in *Lrrk2* KD SIMA9 microglial cells compared to SCR control (**Fig S3B**). Given that *Lrrk2* KO microglial mitochondria had increased membrane potential even under high stress conditions, we next used the Agilent Seahorse Metabolic Analyzer to further investigate the impact of stress on the mitochondrial metabolic state. In the Seahorse assay, oxidative phosphorylation (OXPHOS) and glycolysis are assayed by oxygen consumption rate (OCR) and extracellular acidification rate (ECAR), respectively. We found that OCR in *Lrrk2* KO microglia was enhanced in terms of basal, spare and maximal capacity (**Fig. 3C**, lower quantification), indicating that *Lrrk2* KO mitochondria were not only more active at rest, but they had a greater capacity for maintaining high levels of OXPHOS during stress, indictive of a protected state. This protection was maintained following activation with type I IFN, which can enhance OCR in macrophages (53) (**Fig 3D**, lower quantification), as well as LPS, which has been shown to reduce OCR in macrophages (54) (**Fig 3E**, lower quantification). While oxidative phosphorylation relied heavily on LRRK2, glycolysis, as measured by ECAR, was not impacted under any of these conditions (**Fig S3C-E**). These data are consistent with the mitochondria of *Lrrk2* KO microglia having enhanced protection at the level of mitochondrial homeostasis and suggest that LRRK2 negatively regulates NRF2 to restrict OXPHOS capacity in microglia.

**Figure 3:**
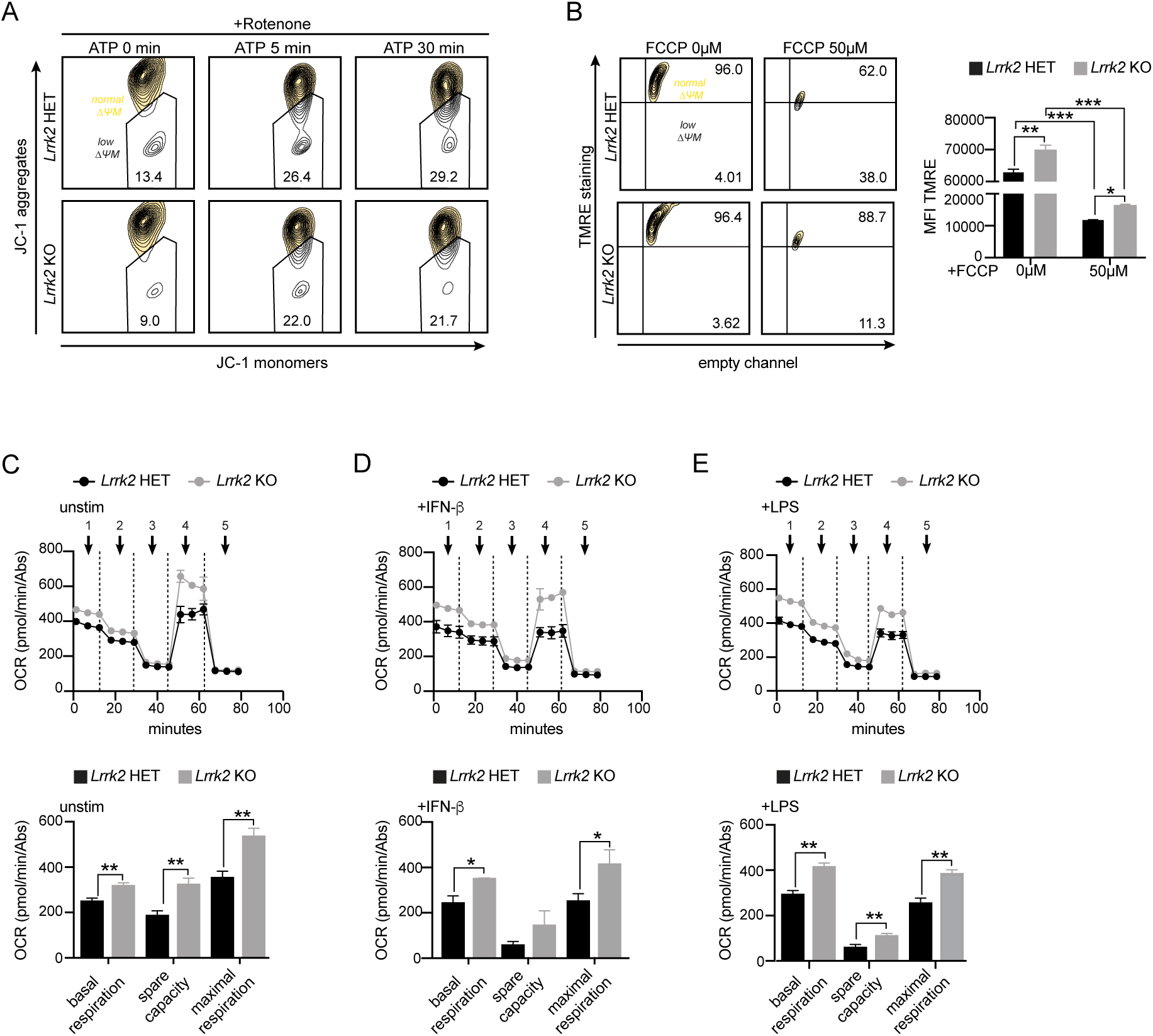
Loss of LRRK2 promotes mitochondrial protection in microglial cells. (**A.**) JC-1 staining of mitochondrial membrane potential in *Lrrk2* KO and HET microglia measured by flow cytometry. Cells were treated with 2.5 µM rotenone for 3 hrs. followed by 5 µM ATP for 5 and 30 min. (**B.**) TMRE staining of mitochondrial membrane potential in *Lrrk2* KO and HET microglia measured by flow cytometry. Cells were treated with vehicle or 50 µM FCCP for 30 min. (**C.**) Oxygen consumption rate (OCR) of resting *Lrrk2* KO and HET microglia measured by the seahorse analyzer mito-stress test. Arrows and numbers indicate reads between injections times. Quantification of major OCR readouts below. (**D.**) The same as in C but cells were treated for 16hrs with 100 IU IFNβ. (**E.**) The same as in (C) and (D) but cells were treated for 16hrs with 10 ng/mL LPS. Two-tailed Student’s t-test or One way ANOVA with Sidak’s multiple comparisons was used to determine statistical significance. *p<0.05, **p<0.01, ***p<0.005.

### Upregulation of NRF2 drives the protection of *Lrrk2* KO microglial cells

In addition to downregulating the type I IFN response directly, NRF2 has been also shown to protect the mitochondria through the upregulation of reactive oxygen scavengers, induction of autophagy, and other protective metabolic programs (55–58). We therefore choose to further explore the divergence between NRF2 in microglial cells and macrophages focusing on NRF2 protective capacity and the mitochondria. To begin to understand how NRF2 might be differentially regulated in *Lrrk2* KO microglial cells compared to controls, we examined both NRF2 expression and localization. Cytoplasmic NRF2 is bound to the KEAP1 complex where it is ubiquitinated and constitutively degraded by the proteosome (59). During cellular stress, KEAP1 releases NRF2, allowing it to rapidly translocate to the nucleus and turn on protective gene expression programs. Consistent with a more activated (protective) state in *Lrrk2* KO microglia, we identified elevated levels of nuclear NRF2 (green) overlapping with DAPI (blue) (**Fig 4A**) and an increase in the overall expression of NRF2. Consistent with enhanced protection we also saw greater induction of nuclear NRF2 in *Lrrk2* KO microglia treated with rotenone for 4h to induce mitochondrial stress (**Fig 4B** right graph). Because *Lrrk2* KO microglia showed enhanced mitochondrial OXPHOS at baseline and during stress with greater OXPHOS reserves, we wanted to determine if that protection was dependent on NRF2. We therefore performed the Seahorse Mito Stress test on *Lrrk2* KO and HET microglia in the presence or absence of NRF2 inhibitors ML385 and brusatol, which inhibit NRF2’s DNA binding ability and expression, respectively (60, 61). Compared to untreaded *Lrrk2* KO microglia (**Fig 4C**, left panel), we found that *Lrrk2* KO microglia treated with ML385 (**Fig 4C**, central panel) or brusatol (**Fig 4C** right panel), lost their protective metabolic state, instead exhibiting reduced basal, spare and maximal respiration potential (**Fig 4D**). Taken together, these data indicate that NRF2 is elevated and activated in *Lrrk2* KO microglia compared to HETs, and this enhanced activity provides protection to the mitochondria.

**Figure 4:**
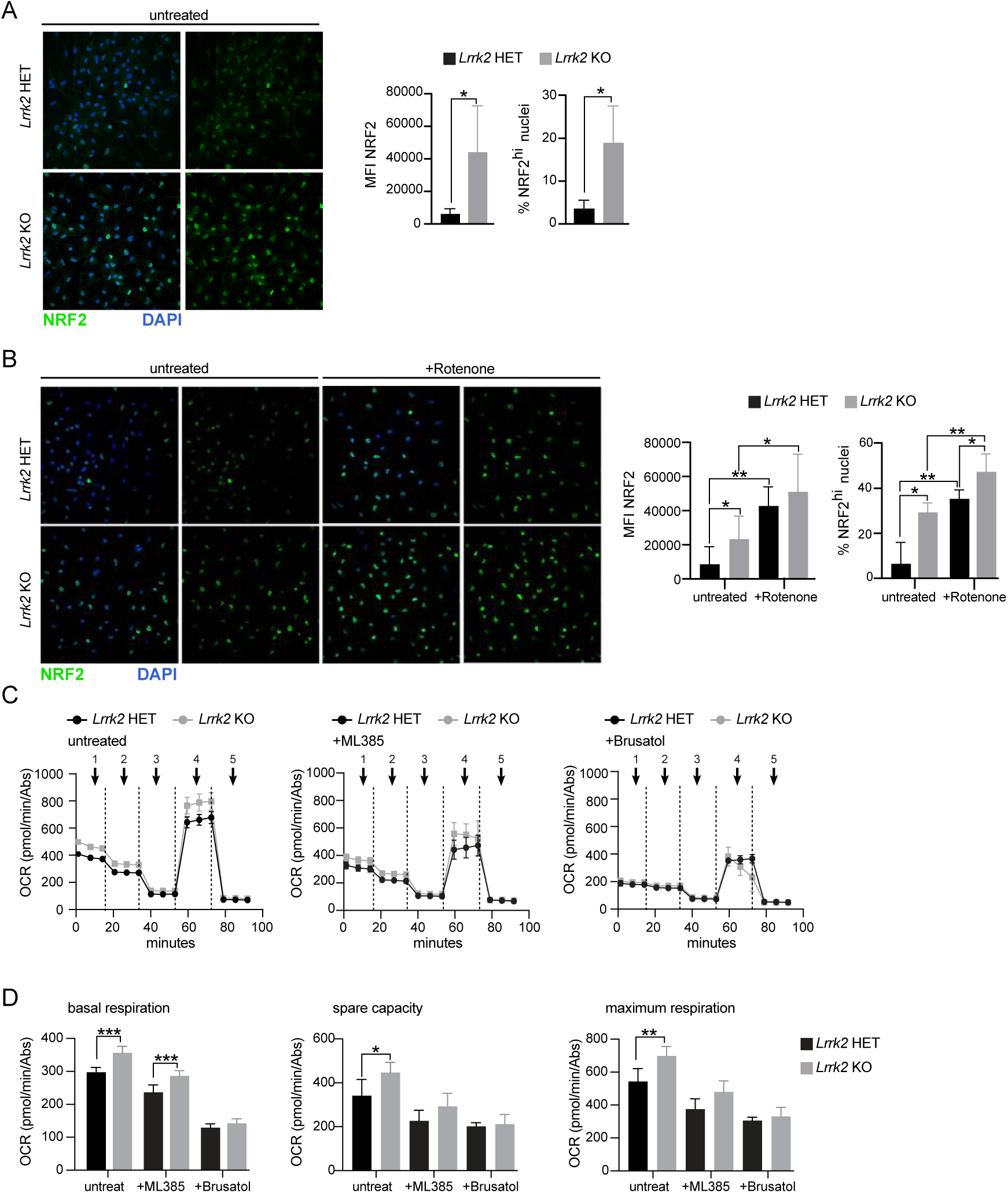
Upregulation of NRF2 drives the mitochondrial protection in LRRK2 KO microglial cells. (**A.**) Protein levels and localization of NRF2 in resting *Lrrk2* KO and HET microglia measured by Immunofluorescence microscopy NRF2 (green), DAPI (blue). left graph measures NRF2 expression based on mean fluorescence intensity, right graph measures number of NRF2^hi^ nuclei per field of vision (**B.**) The same as in (A) but cells were treated with 200 ng/mL rotenone for either 0 or 4 hrs. to induce mitochondrial stress. (**C.**) Oxygen consumption rate (OCR), a proxy for oxidative phosphorylation, of *Lrrk2* KO and HET microglia measured by the seahorse bioanalyzer mito-stress test. Cells were either unstimulated (left panel) or treated with NRF2 inhibitors for 4 hrs., ML385 5 µM (middle panel), Brustol 5 µM (right panel). (**D.**) Quantification of the major OCR readouts from the above line graphs. Two-tailed Student’s t-test or One way ANOVA with Sidak’s multiple comparisons was used to determine statistical significance. *p<0.05, **p<0.01, ***p<0.005.

### LRRK2 kinase activity promotes tonic ISG expression and tempers NRF2 activity

Because increased LRRK2 kinase activity is associated with pathophysiology of both familial and sporadic PD (38, 62, 63), inhibition of LRRK2 kinase activity has been proposed as a possible route of PD intervention. Currently multiple LRRK2 kinase inhibitors are being tested in clinical trials for PD therapeutics (64–66). Given the importance of the type I IFN response to multiple neurodegenerative disorders, we wanted to determine if LRRK2 kinase activity was necessary for controlling ISG expression and restricting NRF2 activity. To this end, we treated SIMA9 and BV2 cells with the blood brain barrier (BBB) penetrant small molecule inhibitor of LRRK2, GNE9605. We found that as early as 24h post-treatment with GNE9605, the ISG transcript *Rsad2* was depleted in both microglial cell lines (**Fig 5A**) and RSAD2 protein levels were reduced in GNE9605-treated SIMA9 cells (**Fig 5B**). LRRK2 kinase activity was also shown to restrict the NRF2 antioxidant response, as SIMA9 microglia had elevated expression of NRF2 and its downstream target HMOX1 (**Fig 5C**) 72h post-treatment with GNE9605. Taken together, these data demonstrate that microglial cells rely on a novel LRRK2-NRF2 circuit to control type I IFN expression in the CNS.

**Figure 5:**
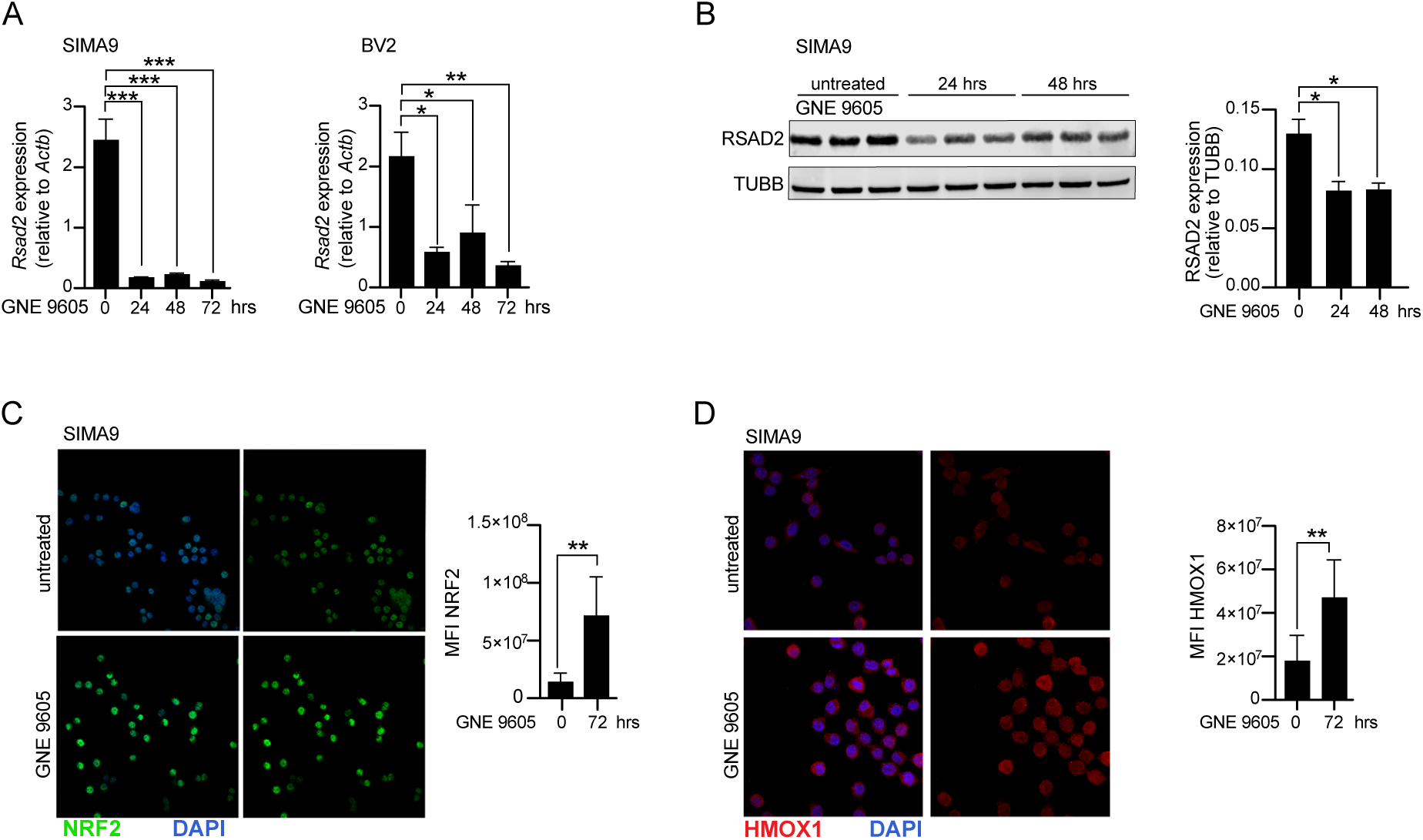
LRRK2 kinase activity regulates the type I IFN response and NRF2 activity in microglial cells. (**A.**) Transcript levels of *Rsad2*, in SIMA9 and BV2 microglia measured by qRT-PCR. Cells were treated for 0, 24, 48, and 72 hrs. with the small molecule LRRK2 inhibitor 10 µM GNE9605. (**B.**) Protein levels of RSAD2, compared to TUBB in SIMA9 microglia measured by western blot. Cells were treated for 24 and 48 hrs. with 10 µM GNE9605 (**C.**) Protein levels and localization of NRF2 in SIMA9 microglia treated with or without 10 µM GNE9605 for 72 hrs. measured by immunofluorescence microscopy NRF2 (green), DAPI (blue). (**D.**) the same as in (C) but looking at HMOX1 (red). Two-tailed Student’s t-test or One way ANOVA with Sidak’s multiple comparisons was used to determine statistical significance. *p<0.05, **p<0.01, ***p<0.005.

## DISCUSSION

Despite being most famously associated with antiviral immunity, the type I IFN response is also necessary for the development of healthy neurons, neuron survival, and neurite outgrowth (17). It follows then, that glial support cells of the CNS need ways to regulate expression of type I IFN and ISGs, both at rest and in response to pathogens or other inflammatory triggers. Here, we found that the PD-associated protein LRRK2 plays a role in regulating the expression of ISGs in microglial cells through restriction of NRF2, a redox factor shown to negatively regulate the type I IFN response in peripheral macrophages. The current macrophage paradigm is that cells are primed via cytosolic sensing of mitochondrial DNA through a cGAS/STING dependent axis to maintain the type I IFN response at a steady state where IFNβ and ISGs can be readily upregulated during infection (67). The capacity of mitochondrial DNA to prime through cGAS/STING has also been shown in microglia from old mice (22). On the contrary, our work shows that another level of type I IFN regulation is present across multiple microglial populations from young mice where a loss of LRRK2 is sufficient to downregulate ISGs and protect mitochondria through upregulation of the redox associated transcription factor NRF2. This suggests that in microglial cells, priming steps are either unnecessary or have detrimental consequences. Given the importance of type I IFN in neuron development, it is understandable that additional regulatory nodes would be necessary to prevent runaway neuroinflammation, which is seen in young mice lacking NRF2 (35, 68–71). The idea that cGAS/STING priming of old microglia is especially intriguing and suggests that this protection is lost over time, possibly to aid in neuroprotection from viruses. At the same time loss of NRF2 protection could also act as a major factor in the pathobiology of diseases like Alzheimer’s and PD which have a strong type I IFN component to disease progression.

In addition to acting as a protective factor in neurodegenerative disease, activation of NRF2 also ameliorates damage caused from ischemic stroke where it protects the BBB, improves edema, and mitigates neurological defects by reducing oxidative stress (69, 72). It is also likely that NRF2 activation restricts the overactivation of type I IFN by microglia which occurs during the acute phase of stroke with deleterious consequences (73). Upregulation of NRF2 could also be exerting protective effects by protecting mitochondria and preventing mitochondrial DNA release where it acts as a danger associated molecular pattern (DAMP) to activate the type IFN response. It is still unclear exactly how loss of LRRK2 leads to the upregulation of NRF2. Better understanding the precise nodes of NRF2 regulation by LRRK2 will provide clarity and insight into the pathology of multiple neurological diseases.

This study shows that LRRK2 can have opposing effects in different macrophage populations. Similarly, LRRK2 has been previously associated with cell-type specific functions in different neuron populations (74, 75). The capacity of LRRK2 to function cell-type specifically indicates a need for more exploration into cell type specific knockouts of this multifunctional protein. Beyond LRRK2, further exploring components of the peripheral immune system in distinct cell types of the CNS could give profound insight into CNS specific damage that occurs during neurological disease. For example, better understanding cell-type specific responses of ISGs, many of which are involved in immune cell recruitment during peripheral infection, could help refine our understanding of peripheral immune cell recruitment to the CNS. Combining this with the study of the NRF2 stress protection axis would give insight into the earliest event of neurodegenerative disease prior to neuron dysfunction and loss.

LRRK2 kinase inhibitors are undergoing clinical trials for protection from PD. Given the reduction in type I IFN signatures which have been shown to be elevated specifically in microglial cells in Alzheimer’s disease as well as traumatic brain injury (19, 73, 76). LRRK2 inhibitor therapy could provide a promising support treatment for patients with multiple different IFN-associated neuropathy to prevent or mitigate excessive damage. This is especially promising given the additional upregulation of NRF2 that could have additional therapeutic benefits through upregulation of protective redox response factors.

**Figure S1:**
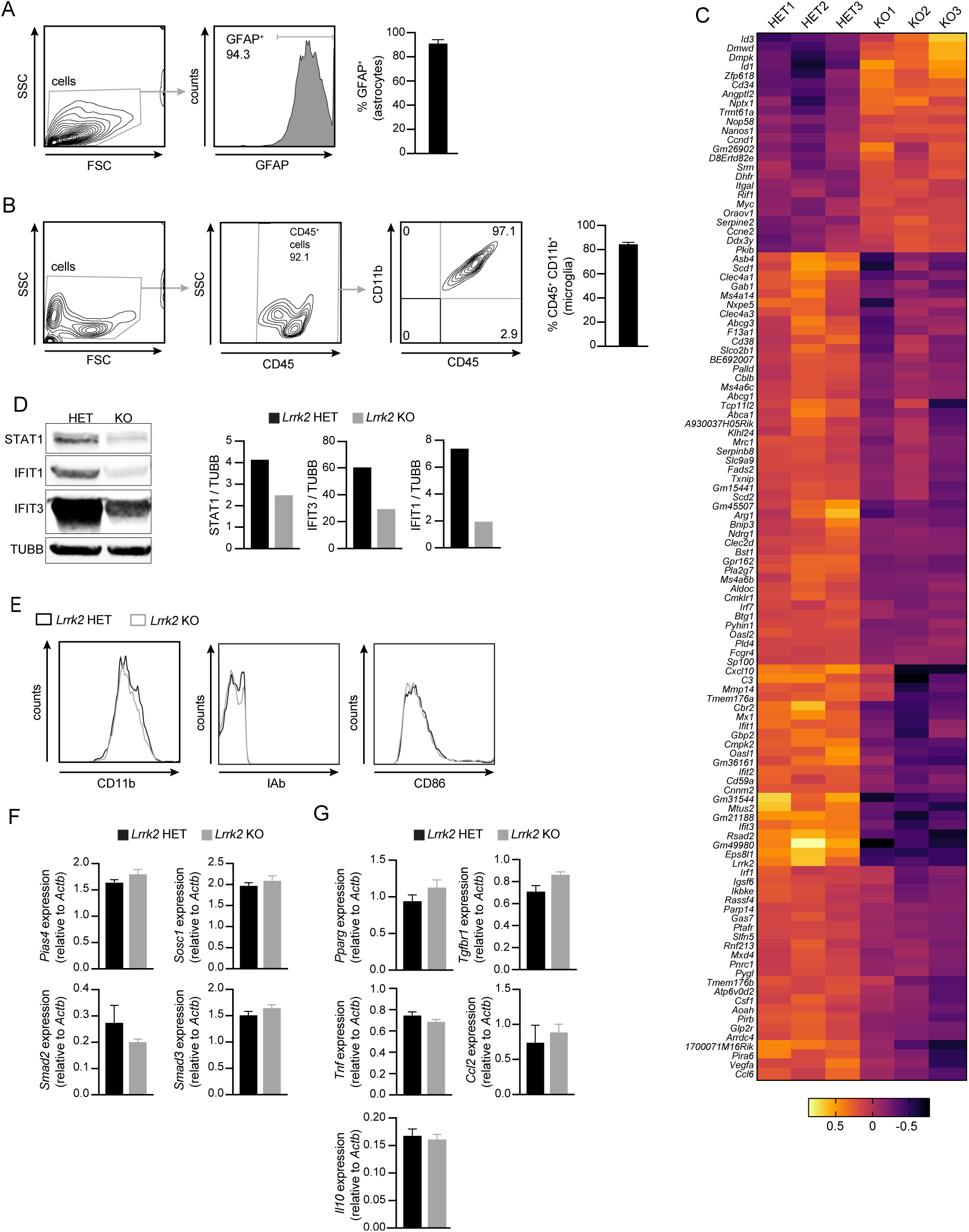
Gating strategy and transcript profile of microglial cells. (**A.**) GFAP expression and cell percentages of astrocyte populations analyzed by flow cytometry. (**B.**) CD45 and CD11b expression and cell percentages of microglial populations analyzed by flow cytometry. (**C.**) Heatmap of significant differentially expressed genes between *Lrrk2* KO microglia and HET controls. (**D.**) Protein levels of ISGs, STAT1, IFIT1, and IFIT3, compared to TUBB in *Lrrk2* KO and HET microglia measured by western blot. (**E.**) CD11b, IAb, CD86 expression of *Lrrk2* KO and HET microglia measured by flow cytometry. (**F.**) Transcript levels of negative regulators of the type I IFN response *Pias4*, *Sosc1*, *Smad2*, *Smad3*, in *Lrrk2* KO and HET microglia measured by qRT-PCR. (**G.**) The same as in (D), but M1 vs M2 macrophage markers *Tnf*, *Ccl2*, *Pparg*, *Tgfbr1*, and *Il10*. Two-tailed Student’s t-test was used to determine statistical significance. *p<0.05, **p<0.01, ***p<0.005.

**Figure S2:**
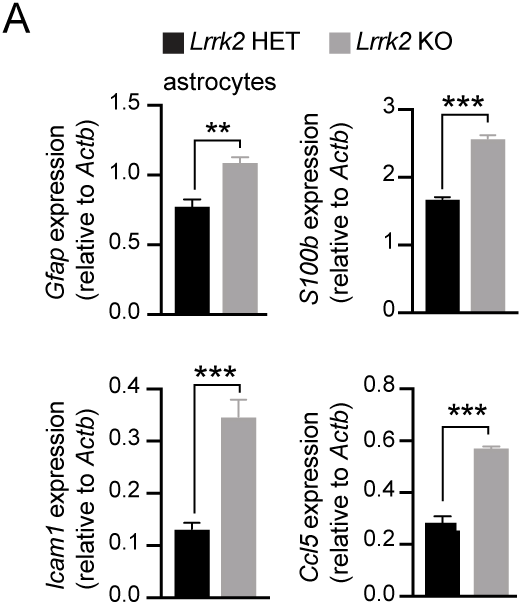
Loss of LRRK2 in astrocytes upregulates activation-associated factors. Transcript levels of astrocyte activation and ISG associated genes *Gfap*, *S100b*, *Icam1*, *Ccl5*, in *Lrrk2* KO and HET astrocytes measured by qRT-PCR. Two-tailed Student’s t-test was used to determine statistical significance. *p<0.05, **p<0.01, ***p<0.005.

**Figure S3:**
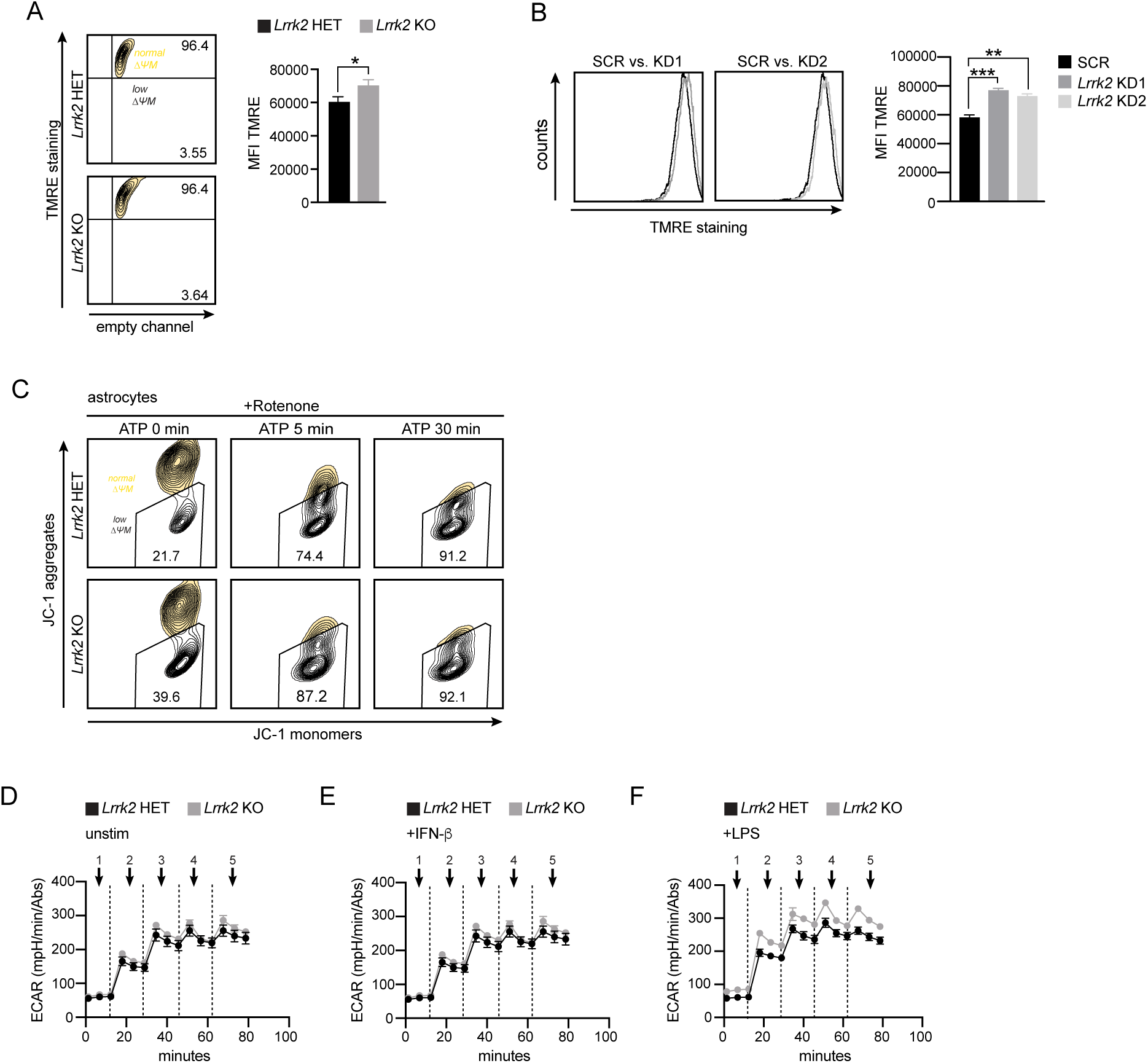
Microglial glycolytic flux is not impacted by a loss of LRRK2. (**A.**) TMRE staining of mitochondrial membrane potential in *Lrrk2* KO and HET microglia measured by flow cytometry. (**B.**) The same as in (A) but cells were SIMA9 microglia *Lrrk2* KD vs. SCR. (**D.**) Extracellular acidification rate (ECAR), a measurement for glycolysis, of unstimulated *Lrrk2* KO and HET microglia (**C.**) JC-1 staining of mitochondrial membrane potential in *Lrrk2* KO and HET astrocytes measured by flow cytometry. Cells were treated with 2.5 µM rotenone for 3 hrs. followed by 5 µM ATP for 5 and 30 min. (**D.**) Extracellular acidification rate (ECAR), a proxy of glycolysis, in resting *Lrrk2* KO and HET microglia measured by the seahorse bioanalyzer mito-stress test. (**E.**) The same as in (D), but cells were treated for 16 hrs. with 100 IU IFNβ (**F.**) The same as in (D) and (E), but cells were treated for 16 hrs. with 10 ng/mL LPS. Two-tailed Student’s t-test or One way ANOVA with Sidak’s multiple comparisons was used to determine statistical significance. *p<0.05, **p<0.01, ***p<0.005.

## MATERIALS AND METHODS

### Mice

*Lrrk2* KO mice (C57BL/6-Lrrk2tm1.1Mjff/J) stock #016121, were purchased from The Jackson Laboratories (Bar Harbor, ME). All mice used in experiments were compared to age- and sex-matched controls by pooling equal males and females between genotypes. To ensure littermate controls were used in all experiments *Lrrk2* KO crosses were made with (KO) *Lrrk2^-/-^* x (HET) *Lrrk2^+/-^* mice. Mice used to make glial cultures were between P1 and P1.5 days old. All animals were housed, bred, and studied at Texas A&M Health Science Center under approved Institutional Care and Use Committee guidelines.

### Primary cells

Mixed glial cultures were differentiated from the brains of neonatal mice as described (41). Briefly, glial cells were isolated from the cortexes of neonate mice at P1-P1.5. Disaggregation media was used to liberate glial cells. Glial cells were centrifuged twice 400 rcf, 5 min and washed in complete media (DMEM, 10% FBS, 1 mM sodium pyruvate, 10% MCSF conditioned media), and grown in 10 mL of media 10 cm TC-treated dishes, one dish per brain at 37 °C 5% CO_2_. Complete media was replaced on day 1 following gentle aspiration. Cells were allowed to differentiate in complete media feeding 5 mL on day 5 and then replacing 5 mL of media on every other day afterward. Following 10 days of culture, microglial cells were isolated from glial cultures by washing briefly with cold 1x PBS + EDTA to detach the microglial layer. Cells were then counted, plated on non-tissue culture treated plates, and washed after 4 hrs with 1x PBS to remove contaminating cell populations.

### Cell lines

SIMA9 cells (ATCC® SC-6004™), were obtained from the ATCC. BV2 cells were gifted by Dr. Jianrong Li, Texas A&M. For BV2 and SIMA9 cells stably expressing scramble knockdown (KD) and *Lrrk2* KD, Lenti-X cells were transfected with a pSICOR scramble non-targeting shRNA construct and pSICOR *Lrrk2* targeting constructs using Polyjet (SignaGen Laboratories). Virus was collected 24 and 48 h post transfection. Microglia were transduced using Lipofectamine 2000 (Thermo Fisher). After 48 h, the media was supplemented with puromycin (Invitrogen) to select cells containing the shRNA plasmid.

### Flow cytometry

To confirm purity, isolated astrocytes were gated by SSC/FSC and identified as GFAP+. Microglial cells isolated from glial cultures were gated on SSC/FSC followed by CD45+ and defined as CD45+ (eBiosciences) CD11b+ (eBiosciences). Activation markers IAb (eBioceiences), and CD86 (eBiosciences) were analyzed on this population. To assess mitochondrial membrane potential, cells were released from culture plates with 1x PBS + EDTA. Single cell suspensions were made in 1x PBS 2% FBS. For TMRE assays, cells were stained for 20 min at 37 °C in 25 nM TMRE dye and analyzed on an LSR Fortessa X20 (BD Biosciences). Fluorescence was measured under PE (585/15). To assess mitochondrial membrane potential under stress, cells were treated for 15 min with 50 µM FCCP. For JC-1 assays, JC-1 dye was sonicated for 5 min with 30 sec intervals. Cells were stained for 20 min at 37 °C in 1 µM JC-1 dye and analyzed on an LSR Fortessa X20 (BD Biosciences). Aggregates were measured under Texas Red (610/20 600 LP) and monomers under FITC (525/50 505 LP). To assess mitochondrial membrane potential under stress, cells were treated for 3 hrs. with 2.5 µM rotenone prior to being lifted of the culture plates. 5 µM ATP was then added for 5, 15, or 30 min, or 50 µM FCCP was added for 15 min.

### Western blot

Cells were lysed in 1x RIPA buffer with protease and phosphatase inhibitors (Pierce). DNA was degraded using 1 U/mL benzonase (EMD Millipore). Proteins were separated by SDS-PAGE and transferred to nitrocellulose membranes. Membranes were blocked overnight in either 5% BSA or non-fat milk, and incubated overnight at 4 °C with the following antibodies: IBA1 (Wako Chemical 019-19741) 1:2000; RSAD2 (Proteintech) 1:1000; STAT1 (Cell Signaling) 1:1000; IFIT1 (Proteintech) 1:1000; IFIT3 (Proteintech) 1:1000; NRF2 (Cell Signaling), 1:1000; HMOX1 (Proteintech); ACTB (Abcam), 1:5000; and TUBB (Abcam), 1:5000. Membranes were incubated with appropriate secondary antibody (Licor) for 2 hrs at RT prior to imaging on Odyssey Fc Dual-Mode Imaging System (Licor).

### Immunofluorescence microscopy

Microglia were seeded at 2.5x10^5^ cells/well on glass coverslips in 24-well dishes. Cells were fixed in 4% paraformaldehyde for 10 min at RT and then washed three times with PBS. Coverslips were incubated in primary antibody diluted in 1x PBS + 5% non-fat milk + 0.1% Triton-X (PBS-MT) for 3 hrs. Cells were then washed three times in 1x PBS and incubated in secondary antibodies and DAPI diluted in PBS-MT for 1 hrs. Coverslips were washed twice with 1x PBS and twice with deionized water and mounted on glass slides using Prolong Gold Antifade Reagent (Invitrogen)

### Seahorse metabolic assays

Seahorse XF Mito Stress test kits and cartridges (Agilent) were prepared per manufacturers protocol and as previously described (42). Microglia were seeded at 5x10^4^ cells/well and analyzed the following day on a Seahorse XF 96well Analyzer (Agilent). For treatments cells were stimulated overnight with 10 ng/ml LPS (Invivogen), or 100 IU IFN-β (PBL), or for 4 hrs with 5 µM ML385 (Selleckchem) or 5 µM Brusatol (Selleckchem) for NRF2 inhibition.

### mRNA sequencing

Microglial cell library preparation was carried out by the Baylor College of Medicine Genomic and RNA Profiling Core (GARP) in biological triplicate. RNA sequencing (150 bp paired-end reads) was performed on an Illumina NovaSeq 6000 with S4 flow cell. Data was analyzed by ROSALIND® (https://rosalind.bio/), with a HyperScale architecture developed by ROSALIND, Inc. (San Diego, CA). Quality scores were assessed using FastQC. Reads were aligned to the *Mus musculus* genome build GRCm39 using Agilent software. Differentially expressed genes were selected as those with p-value threshold <0.05 in the heatmaps represented. Transcriptome analysis was performed using IPA analysis to generate GO term, disease pathway lists, and to compare loss of LRRK2 between microglia and macrophages. Heatmaps were generated using GraphPad Prism software (GraphPad, San Diego, CA). Rosalind and IPA were used for pathways analysis and to generate volcano plots and Venn diagrams.

### qRT-PCR

RNA was isolated using Directzol RNAeasy kits (Zymogen). cDNA was made with iScript Direct Synthesis kits (BioRad) per manufacturer’s protocol. qRT-PCR was performed in triplicate using Sybr Green Power up (ThermoFisher). Data was analyzed on a ViiA 7 Real-Time PCR System (Applied Biosystems).

### Statistical analysis

All data are representative of 3 or more independent experiments with an n=3 or more. Graphs were generated using Prism (GraphPad). Significance for assays were determined using a student’s two-tailed t-test, or a one-way ANOVA followed by a Sidak’s multiple comparisons test for more than two variables, unless otherwise noted.

## ACKNOWLEDGEMENTS

We would like to acknowledge the members of the Patrick and Watson labs for their helpful discussions and feedback. We would particularly like to thank Haley Scott for help with microglial cell isolation practices and written protocol development. We’d like to thank the Li lab for their help with the culture of primary microglial cells and gift of BV2 cells. We’d also like to thank A. Phillip West and the West lab for their help with mitochondrial experiments.

## AUTHOR CONTRIBUTIONS

Conceptualization, C.G.W., K.L.P., R.O.W.; Investigation, C.G.W., L.M.E., A.J.C.; Methodology, C.G.W. A.J.C., L.M.E.; Writing, C.G.W., L.M.E, K.L.P., R.O.W.; Visualization, C.G.W., K.L.P. and R.O.W.; Funding acquisition, R.O.W., C.G.W.; Supervision, R.O.W., K.L.P.

## REFERENCES

1. Askew, K., K. Li, A. Olmos-Alonso, F. Garcia-Moreno, Y. Liang, P. Richardson, T. Tipton, M. A. Chapman, K. Riecken, S. Beccari, A. Sierra, Z. Molnár, M. S. Cragg, O. Garaschuk, V. H. Perry, and D. Gomez-Nicola. 2017. Coupled Proliferation and Apoptosis Maintain the Rapid Turnover of Microglia in the Adult Brain. Cell Rep 18.

2. Kierdorf, K., D. Erny, T. Goldmann, V. Sander, C. Schulz, E. G. Perdiguero, P. Wieghofer, A. Heinrich, P. Riemke, C. Hölscher, D. N. Müller, B. Luckow, T. Brocker, K. Debowski, G. Fritz, G. Opdenakker, A. Diefenbach, K. Biber, M. Heikenwalder, F. Geissmann, F. Rosenbauer, and M. Prinz. 2013. Microglia emerge from erythromyeloid precursors via Pu.1-and Irf8-dependent pathways. Nat Neurosci 16.

3. Ginhoux, F., S. Lim, G. Hoeffel, D. Low, and T. Huber. 2013. Origin and differentiation of microglia. Front Cell Neurosci.

4. Ginhoux, F., M. Greter, M. Leboeuf, S. Nandi, P. See, S. Gokhan, M. F. Mehler, S. J. Conway, L. G. Ng, E. R. Stanley, I. M. Samokhvalov, and M. Merad. 2010. Fate mapping analysis reveals that adult microglia derive from primitive macrophages. Science (1979) 330.

5. Alliot, F., I. Godin, and B. Pessac. 1999. Microglia derive from progenitors, originating from the yolk sac, and which proliferate in the brain. Developmental Brain Research 117.

6. Gomez Perdiguero, E., K. Klapproth, C. Schulz, K. Busch, E. Azzoni, L. Crozet, H. Garner, C. Trouillet, M. F. De Bruijn, F. Geissmann, and H. R. Rodewald. 2015. Tissue-resident macrophages originate from yolk-sac-derived erythro-myeloid progenitors. Nature 518.

7. Paolicelli, R. C., G. Bolasco, F. Pagani, L. Maggi, M. Scianni, P. Panzanelli, M. Giustetto, T. A. Ferreira, E. Guiducci, L. Dumas, D. Ragozzino, and C. T. Gross. 2011. Synaptic pruning by microglia is necessary for normal brain development. Science (1979) 333.

8. Zengeler, K. E., and J. R. Lukens. 2021. Innate immunity at the crossroads of healthy brain maturation and neurodevelopmental disorders. Nat Rev Immunol 21.

9. Salter, M. W., and B. Stevens. 2017. Microglia emerge as central players in brain disease. Nat Med 23.

10. Chien, C.-H., M.-J. Lee, H.-C. Liou, H.-H. Liou, and W.-M. Fu. 2016. Microglia-Derived Cytokines/Chemokines Are Involved in the Enhancement of LPS-Induced Loss of Nigrostriatal Dopaminergic Neurons in DJ-1 Knockout Mice. PLoS One 11: e0151569.

11. Davalos, D., J. Grutzendler, G. Yang, J. V. Kim, Y. Zuo, S. Jung, D. R. Littman, M. L. Dustin, and W. B. Gan. 2005. ATP mediates rapid microglial response to local brain injury in vivo. Nat Neurosci 8.

12. Cowan, M. N., I. Sethi, and T. H. Harris. 2022. Microglia in CNS infections: insights from Toxoplasma gondii and other pathogens. Trends Parasitol 38.

13. McGeer, P. L., S. Itagaki, H. Akiyama, and E. G. McGeer. 1988. Rate of cell death in parkinsonism indicates active neuropathological process. Ann Neurol 24: 574–576.

14. McGeer, P. L., S. Itagaki, B. E. Boyes, and E. G. McGeer. 1988. Reactive microglia are positive for HLA-DR in the: Substantia nigra of Parkinson’s and Alzheimer’s disease brains. Neurology 38.

15. Bartels, T., S. De Schepper, and S. Hong. 2020. Microglia modulate neurodegeneration in Alzheimer’s and Parkinson’s diseases. Science (1979) 370.

16. Muzio, L., A. Viotti, and G. Martino. 2021. Microglia in Neuroinflammation and Neurodegeneration: From Understanding to Therapy. Front Neurosci 15.

17. Ejlerskov, P., J. G. Hultberg, J. Y. Wang, R. Carlsson, M. Ambjørn, M. Kuss, Y. Liu, G. Porcu, K. Kolkova, C. Friis Rundsten, K. Ruscher, B. Pakkenberg, T. Goldmann, D. Loreth, M. Prinz, D. C. Rubinsztein, and S. Issazadeh-Navikas. 2015. Lack of Neuronal IFN-β-IFNAR Causes Lewy Body- and Parkinson’s Disease-like Dementia. Cell 163.

18. Escoubas, C. C., L. C. Dorman, P. T. Nguyen, C. Lagares-Linares, H. Nakajo, S. R. Anderson, J. J. Barron, S. D. Wade, B. Cuevas, I. D. Vainchtein, N. J. Silva, R. Guajardo, Y. Xiao, P. V. Lidsky, E. Y. Wang, B. M. Rivera, S. E. Taloma, D. K. Kim, E. Kaminskaya, H. Nakao-Inoue, B. Schwer, T. D. Arnold, A. B. Molofsky, C. Condello, R. Andino, T. J. Nowakowski, and A. V. Molofsky. 2024. Type-I-interferon-responsive microglia shape cortical development and behavior. Cell 187.

19. Roy, E. R., B. Wang, Y. W. Wan, G. Chiu, A. Cole, Z. Yin, N. E. Propson, Y. Xu, J. L. Jankowsky, Z. Liu, V. M. Y. Lee, J. Q. Trojanowski, S. D. Ginsberg, O. Butovsky, H. Zheng, and W. Cao. 2020. Type I interferon response drives neuroinflammation and synapse loss in Alzheimer disease. Journal of Clinical Investigation 130.

20. Roy, E. R., G. Chiu, S. Li, N. E. Propson, R. Kanchi, B. Wang, C. Coarfa, H. Zheng, and W. Cao. 2022. Concerted type I interferon signaling in microglia and neural cells promotes memory impairment associated with amyloid β plaques. Immunity 55.

21. Hammond, T. R., C. Dufort, L. Dissing-Olesen, S. Giera, A. Young, A. Wysoker, A. J. Walker, F. Gergits, M. Segel, J. Nemesh, S. E. Marsh, A. Saunders, E. Macosko, F. Ginhoux, J. Chen, R. J. M. Franklin, X. Piao, S. A. McCarroll, and B. Stevens. 2019. Single-Cell RNA Sequencing of Microglia throughout the Mouse Lifespan and in the Injured Brain Reveals Complex Cell-State Changes. Immunity 50.

22. Gulen, M. F., N. Samson, A. Keller, M. Schwabenland, C. Liu, S. Glück, V. V. Thacker, L. Favre, B. Mangeat, L. J. Kroese, P. Krimpenfort, M. Prinz, and A. Ablasser. 2023. cGAS–STING drives ageing-related inflammation and neurodegeneration. Nature 620.

23. Gunderstofte, C., M. B. Iversen, S. Peri, A. Thielke, S. Balachandran, C. K. Holm, and D. Olagnier. 2019. Nrf2 Negatively Regulates Type I Interferon Responses and Increases Susceptibility to Herpes Genital Infection in Mice. Front Immunol 10.

24. Wyler, E., V. Franke, J. Menegatti, C. Kocks, A. Boltengagen, S. Praktiknjo, B. Walch-Rückheim, J. Bosse, N. Rajewsky, F. Grässer, A. Akalin, and M. Landthaler. 2019. Single-cell RNA-sequencing of herpes simplex virus 1-infected cells connects NRF2 activation to an antiviral program. Nat Commun 10.

25. Ryan, D. G., E. V. Knatko, A. M. Casey, J. L. Hukelmann, S. Dayalan Naidu, A. J. Brenes, T. Ekkunagul, C. Baker, M. Higgins, L. Tronci, E. Nikitopolou, T. Honda, R. C. Hartley, L. A. J. O’Neill, C. Frezza, A. I. Lamond, A. Y. Abramov, J. S. C. Arthur, D. A. Cantrell, M. P. Murphy, and A. T. Dinkova-Kostova. 2022. Nrf2 activation reprograms macrophage intermediary metabolism and suppresses the type I interferon response. iScience 25.

26. Thimmulappa, R. K., H. Lee, T. Rangasamy, S. P. Reddy, M. Yamamoto, T. W. Kensler, and S. Biswal. 2006. Nrf2 is a critical regulator of the innate immune response and survival during experimental sepsis. Journal of Clinical Investigation 116.

27. Kobayashi, E. H., T. Suzuki, R. Funayama, T. Nagashima, M. Hayashi, H. Sekine, N. Tanaka, T. Moriguchi, H. Motohashi, K. Nakayama, and M. Yamamoto. 2016. Nrf2 suppresses macrophage inflammatory response by blocking proinflammatory cytokine transcription. Nat Commun 7.

28. Olagnier, D., S. Peri, C. Steel, N. van Montfoort, C. Chiang, V. Beljanski, M. Slifker, Z. He, C. N. Nichols, R. Lin, S. Balachandran, and J. Hiscott. 2014. Cellular Oxidative Stress Response Controls the Antiviral and Apoptotic Programs in Dengue Virus-Infected Dendritic Cells. PLoS Pathog 10.

29. Olagnier, D., R. R. Lababidi, S. B. Hadj, A. Sze, Y. Liu, S. D. Naidu, M. Ferrari, Y. Jiang, C. Chiang, V. Beljanski, M. L. Goulet, E. V. Knatko, A. T. Dinkova-Kostova, J. Hiscott, and R. Lin. 2017. Activation of Nrf2 Signaling Augments Vesicular Stomatitis Virus Oncolysis via Autophagy-Driven Suppression of Antiviral Immunity. Molecular Therapy 25.

30. Olagnier, D., E. Farahani, J. Thyrsted, J. Blay-Cadanet, A. Herengt, M. Idorn, A. Hait, B. Hernaez, A. Knudsen, M. B. Iversen, M. Schilling, S. E. Jørgensen, M. Thomsen, L. S. Reinert, M. Lappe, H. D. Hoang, V. H. Gilchrist, A. L. Hansen, R. Ottosen, C. G. Nielsen, C. Møller, D. van der Horst, S. Peri, S. Balachandran, J. Huang, M. Jakobsen, E. B. Svenningsen, T. B. Poulsen, L. Bartsch, A. L. Thielke, Y. Luo, T. Alain, J. Rehwinkel, A. Alcamí, J. Hiscott, T. Mogensen, S. R. Paludan, and C. K. Holm. 2020. SARS-CoV2-mediated suppression of NRF2-signaling reveals potent antiviral and anti-inflammatory activity of 4-octyl-itaconate and dimethyl fumarate. Nat Commun 11.

31. Olagnier, D., A. M. Brandtoft, C. Gunderstofte, N. L. Villadsen, C. Krapp, A. L. Thielke, A. Laustsen, S. Peri, A. L. Hansen, L. Bonefeld, J. Thyrsted, V. Bruun, M. B. Iversen, L. Lin, V. M. Artegoitia, C. Su, L. Yang, R. Lin, S. Balachandran, Y. Luo, M. Nyegaard, B. Marrero, R. Goldbach-Mansky, M. Motwani, D. G. Ryan, K. A. Fitzgerald, L. A. O’Neill, A. K. Hollensen, C. K. Damgaard, F. v. de Paoli, H. C. Bertram, M. R. Jakobsen, T. B. Poulsen, and C. K. Holm. 2018. Nrf2 negatively regulates STING indicating a link between antiviral sensing and metabolic reprogramming. Nat Commun 9.

32. Wu, M., Y. Fan, L. Li, and J. Yuan. 2024. Bi-directional regulation of type I interferon signaling by heme oxygenase-1. iScience 27.

33. Hubbs, A. F., S. A. Benkovic, D. B. Miller, J. P. O’Callaghan, L. Battelli, D. Schwegler-Berry, and M. Qiang. 2007. Vacuolar leukoencephalopathy with widespread astrogliosis in mice lacking transcription factor Nrf2. American Journal of Pathology 170.

34. Jazwa, A., A. I. Rojo, N. G. Innamorato, M. Hesse, J. Fernández-Ruiz, and A. Cuadrado. 2011. Pharmacological targeting of the transcription factor NRf2 at the basal ganglia provides disease modifying therapy for experimental parkinsonism. Antioxid Redox Signal 14.

35. Rojo, A. I., N. G. Innamorato, A. M. Martín-Moreno, M. L. De Ceballos, M. Yamamoto, and A. Cuadrado. 2010. Nrf2 regulates microglial dynamics and neuroinflammation in experimental Parkinson’s disease. Glia 58.

36. Giesert, F., A. Hofmann, A. Bürger, J. Zerle, K. Kloos, U. Hafen, L. Ernst, J. Zhang, D. M. Vogt-Weisenhorn, and W. Wurst. 2013. Expression Analysis of Lrrk1, Lrrk2 and Lrrk2 Splice Variants in Mice. PLoS One 8: e63778.

37. Rocha, E. M., M. T. Keeney, R. Di Maio, B. R. De Miranda, and J. T. Greenamyre. 2022. LRRK2 and idiopathic Parkinson’s disease. Trends Neurosci 45: 224–236.

38. Kluss, J. H., A. Mamais, and M. R. Cookson. 2019. LRRK2 links genetic and sporadic Parkinson’s disease. Biochem. Soc. Trans. 47: 651–661.

39. Smith, W. W., Z. Pei, H. Jiang, V. L. Dawson, T. M. Dawson, and C. A. Ross. 2006. Kinase activity of mutant LRRK2 mediates neuronal toxicity. Nat Neurosci 9: 1231–1233.

40. Weindel, C. G., S. L. Bell, K. J. Vail, K. O. West, K. L. Patrick, and R. O. Watson. 2020. LRRK2 maintains mitochondrial homeostasis and regulates innate immune responses to Mycobacterium tuberculosis. Elife 9.

41. Lian, H., E. Roy, and H. Zheng. 2016. Protocol for Primary Microglial Culture Preparation. Bio Protoc 6.

42. Van den Bossche, J., J. Baardman, and M. P. J. de Winther. 2015. Metabolic characterization of polarized M1 and M2 bone marrow-derived macrophages using real-time extracellular flux analysis. Journal of Visualized Experiments 2015.

43. Quan, Y., J. Xu, Q. Xu, Z. Guo, R. Ou, H. Shang, and Q. Wei. 2023. Association between the risk and severity of Parkinson’s disease and plasma homocysteine, vitamin B12 and folate levels: a systematic review and meta-analysis. Front Aging Neurosci 15.

44. Krawczyk, M. C., M. Godoy, P. Vander, A. J. Zhang, and Y. Zhang. 2023. Loss of Serpin E2 alters antimicrobial gene expression by microglia but not astrocytes. Neurosci Lett 811.

45. Asheuer, M., F. Pflumio, S. Benhamida, A. Dubart-Kupperschmitt, F. Fouquet, Y. Imai, P. Aubourg, and N. Cartier. 2004. Human CD34+ cells differentiate into microglia and express recombinant therapeutic protein. Proc Natl Acad Sci U S A 101.

46. Caillet-Boudin, M. L., F. J. Fernandez-Gomez, H. Tran, C. M. Dhaenens, L. Buee, and N. Sergeant. 2014. Brain pathology in myotonic dystrophy: When tauopathy meets spliceopathy and RNAopathy. Front Mol Neurosci 6.

47. Tzeng, S. F., and J. De Vellis. 1998. Id1, Id2, and Id3 gene expression in neural cells during development. Glia 24.

48. Lee, A. J., and A. A. Ashkar. 2018. The dual nature of type I and type II interferons. Front Immunol 9.

49. Ivashkiv, L. B., and L. T. Donlin. 2014. Regulation of type i interferon responses. Nat Rev Immunol 14: 36–49.

50. Arimoto, K. I., S. Miyauchi, S. A. Stoner, J. B. Fan, and D. E. Zhang. 2018. Negative regulation of type I IFN signaling. J Leukoc Biol 103.

51. Tanaka, T., K. Murakami, Y. Bando, and S. Yoshida. 2015. Interferon regulatory factor 7 participates in the M1-like microglial polarization switch. Glia 63.

52. Liddelow, S. A., K. A. Guttenplan, L. E. Clarke, F. C. Bennett, C. J. Bohlen, L. Schirmer, M. L. Bennett, A. E. Münch, W.-S. Chung, T. C. Peterson, D. K. Wilton, A. Frouin, B. A. Napier, N. Panicker, M. Kumar, M. S. Buckwalter, D. H. Rowitch, V. L. Dawson, T. M. Dawson, B. Stevens, and B. A. Barres. 2017. Neurotoxic reactive astrocytes are induced by activated microglia. Nature 541: 481–487.

53. Wu, D., D. E. Sanin, B. Everts, Q. Chen, J. Qiu, M. D. Buck, A. Patterson, A. M. Smith, C. H. Chang, Z. Liu, M. N. Artyomov, E. L. Pearce, M. Cella, and E. J. Pearce. 2016. Type 1 Interferons Induce Changes in Core Metabolism that Are Critical for Immune Function. Immunity 44.

54. Kelly, B., and L. A. O’Neill. 2015. Metabolic reprogramming in macrophages and dendritic cells in innate immunity. Cell Res 25: 771–784.

55. Abdalkader, M., R. Lampinen, K. M. Kanninen, T. M. Malm, and J. R. Liddell. 2018. Targeting Nrf2 to suppress ferroptosis and mitochondrial dysfunction in neurodegeneration. Front Neurosci 12.

56. Gumeni, S., E. D. Papanagnou, M. S. Manola, and I. P. Trougakos. 2021. Nrf2 activation induces mitophagy and reverses Parkin/Pink1 knock down-mediated neuronal and muscle degeneration phenotypes. Cell Death Dis 12.

57. Bento-Pereira, C., and A. T. Dinkova-Kostova. 2021. Activation of transcription factor Nrf2 to counteract mitochondrial dysfunction in Parkinson’s disease. Med Res Rev 41.

58. Cvetko, F., S. T. Caldwell, M. Higgins, T. Suzuki, M. Yamamoto, H. A. Prag, R. C. Hartley, A. T. Dinkova-Kostova, and M. P. Murphy. 2021. Nrf2 is activated by disruption of mitochondrial thiol homeostasis but not by enhanced mitochondrial superoxide production. Journal of Biological Chemistry 296.

59. Baird, L., and M. Yamamoto. 2020. The Molecular Mechanisms Regulating the KEAP1-NRF2 Pathway. Mol Cell Biol 40.

60. Singh, A., S. Venkannagari, K. H. Oh, Y. Q. Zhang, J. M. Rohde, L. Liu, S. Nimmagadda, K. Sudini, K. R. Brimacombe, S. Gajghate, J. Ma, A. Wang, X. Xu, S. A. Shahane, M. Xia, J. Woo, G. A. Mensah, Z. Wang, M. Ferrer, E. Gabrielson, Z. Li, F. Rastinejad, M. Shen, M. B. Boxer, and S. Biswal. 2016. Small molecule inhibitor of NRF2 selectively intervenes therapeutic resistance in KEAP1-deficient NSCLC tumors. ACS Chem Biol 11.

61. Olayanju, A., I. M. Copple, H. K. Bryan, G. T. Edge, R. L. Sison, M. W. Wong, Z. Q. Lai, Z. X. Lin, K. Dunn, C. M. Sanderson, A. F. Alghanem, M. J. Cross, E. C. Ellis, M. Ingelman-Sundberg, H. Z. Malik, N. R. Kitteringham, C. E. Goldring, and B. K. Park. 2015. Brusatol provokes a rapid and transient inhibition of Nrf2 signaling and sensitizes mammalian cells to chemical toxicity-Implications for therapeutic targeting of Nrf2. Free Radic Biol Med 78.

62. Ilieva, N. M., E. K. Hoffman, M. A. Ghalib, J. T. Greenamyre, and B. R. De Miranda. 2024. LRRK2 kinase inhibition protects against Parkinson’s disease-associated environmental toxicants. Neurobiol Dis 196.

63. Lubben, N., J. K. Brynildsen, C. M. Webb, H. L. Li, C. E. G. Leyns, L. Changolkar, B. Zhang, E. S. Meymand, M. O’Reilly, Z. Madaj, D. DeWeerd, M. J. Fell, V. M. Y. Lee, D. S. Bassett, and M. X. Henderson. 2024. LRRK2 kinase inhibition reverses G2019S mutation-dependent effects on tau pathology progression. Transl Neurodegener 13.

64. West, A. B. 2017. Achieving neuroprotection with LRRK2 kinase inhibitors in Parkinson disease. Exp Neurol 298.

65. Wojewska, D. N., and A. Kortholt. 2021. Lrrk2 targeting strategies as potential treatment of parkinson’s disease. Biomolecules 11.

66. Taymans, J. M., M. Fell, T. Greenamyre, W. D. Hirst, A. Mamais, S. Padmanabhan, I. Peter, H. Rideout, and A. Thaler. 2023. Perspective on the current state of the LRRK2 field. NPJ Parkinsons Dis 9.

67. West, A. P., W. Khoury-Hanold, M. Staron, M. C. Tal, C. M. Pineda, S. M. Lang, M. Bestwick, B. A. Duguay, N. Raimundo, D. A. Macduff, S. M. Kaech, J. R. Smiley, R. E. Means, A. Iwasaki, and G. S. Shadel. Mitochondrial DNA stress primes the antiviral innate immune response.

68. Shah, Z. A., R. C. Li, R. K. Thimmulappa, T. W. Kensler, M. Yamamoto, S. Biswal, and S. Doré. 2007. Role of reactive oxygen species in modulation of Nrf2 following ischemic reperfusion injury. Neuroscience 147.

69. Shih, A. Y., P. Li, and T. H. Murphy. 2005. A small-molecule-inducible Nrf2-mediated antioxidant response provides effective prophylaxis against cerebral ischemia in vivo. Journal of Neuroscience 25.

70. Chen, P. C., M. R. Vargas, A. K. Pani, R. J. Smeyne, D. A. Johnson, Y. W. Kan, and J. A. Johnson. 2009. Nrf2-mediated neuroprotection in the MPTP mouse model of Parkinson’s disease: Critical role for the astrocyte. Proc Natl Acad Sci U S A 106.

71. Branca, C., E. Ferreira, T. V. Nguyen, K. Doyle, A. Caccamo, and S. Oddo. 2017. Genetic reduction of Nrf2 exacerbates cognitive deficits in a mouse model of Alzheimer’s disease. Hum Mol Genet 26.

72. Fan, W., H. Chen, M. Li, X. Fan, F. Jiang, C. Xu, Y. Wang, W. Wei, J. Song, D. Zhong, and G. Li. 2024. NRF2 activation ameliorates blood–brain barrier injury after cerebral ischemic stroke by regulating ferroptosis and inflammation. Sci Rep 14.

73. Cui, P., B. Song, Z. Xia, and Y. Xu. 2024. Type I Interferon Signalling and Ischemic Stroke: Mechanisms and Therapeutic Potentials. Transl Stroke Res.

74. Zhao, Y., N. Vavouraki, R. C. Lovering, V. Escott-Price, K. Harvey, P. A. Lewis, and C. Manzoni. 2023. Tissue specific LRRK2 interactomes reveal a distinct striatal functional unit. PLoS Comput Biol 19.

75. Skelton, P. D., V. Tokars, and L. Parisiadou. 2022. LRRK2 at Striatal Synapses: Cell-Type Specificity and Mechanistic Insights. Cells 11.

76. Jassam, Y. N., S. Izzy, M. Whalen, D. B. McGavern, and J. El Khoury. 2017. Neuroimmunology of Traumatic Brain Injury: Time for a Paradigm Shift. Neuron 95.

